# Capacity building needed to reap the benefits of access to biodiversity collections

**DOI:** 10.1101/2024.11.12.623213

**Authors:** Quentin Groom, Sofie Meeus, Sara Barrios, Catherine Childs, Colin Clubbe, Ernestine Corbett, Sarita Francis, Alan Gray, Luke Harding, Annick Jackman, Rebecca Machin, Andrew McGovern, Mike Pienkowski, Delmaude Ryan, Chris Sealys, Catherine Wensink, Jodey Peyton

## Abstract

**Summary:**

1. This research examines biodiversity specimens from two areas of the Caribbean to understand patterns of collection and the roles of the people involved. Using open data from the Global Biodiversity Information Facility (GBIF) and Wikidata, we aimed to uncover geographic and historical trends in specimen use. This study aims to provide concrete evidence to guide collaboration between collection-holding institutions and the communities that need their resources most.
2. We analysed biodiversity specimens from Montserrat and the Cayman Islands in three steps. First, we extracted specimen data from GBIF, disambiguated collector names, and linked them to unique biographical entries. Next, we connected collectors to their publications and specimens. Finally, we analysed the modern use of these specimens through citation data, mapping author affiliations and research themes.
3. Specimens are predominantly housed in the Global North and were initially used by their collectors, whose focus was largely on taxonomy and biogeography. With digitisation, use of these collections remains concentrated in the Global North and covers a broader range of subjects, although Brazil and China stand out as significant users of digital collection data compared to other similar countries.
4. The availability of open digital data from collections in the Global North has led to a substantial increase in the reuse of these data across biodiversity science. Nonetheless, most research using these data is still conducted in the Global North. For the non-monetary benefits of digitisation to extend to the countries of origin, capacity building in the Global South is crucial, Open Data alone are insufficient.

**Societal Impact Statement:** Digital biodiversity data from herbaria and museums hold significant potential for nature conservation in the Global South, yet many regions, like Montserrat and the Cayman Islands in the Caribbean, are, for multiple reasons, unable to fully leverage this information. This lack of skills and resources limits local conservation efforts, showing the need for more investment in training, facilities, and expertise. Although past funding has helped improve coordination and build skills, our findings show that more work is needed to make sure conservation in these biodiverse areas can continue in the long term.

## Introduction

Museums and herbaria are becoming increasingly more open with their natural history collections. They are improving access to collections digitally (Drew et al., 2017); and conducting research in a more collaborative and co-creational way (Ariese & Wróblewska, 2021); They are also being more transparent about the colonial history of scientific discovery (Narkiss, 2022; Park et al., 2023; Wintle, 2016).

Nevertheless, much change is still required to reverse the influence of colonial history and the subject remains controversial (Maranda, 2021). Unequal access is seen, particularly in the Global South, as contributing to biopiracy of resources from biodiverse countries, including through “parachute science” and “scientific tourism”, where researchers from the Global North conduct research and extract specimens from countries in the Global South without any involvement of, or concern for local communities (Stefanoudis et al., 2021).

The inequality of access to data was acknowledged by the Convention on Biological Diversity, resulting in the establishment of the Nagoya Protocol. This protocol aims to ensure the fair sharing of benefits derived from biodiversity (Secretariat of the Convention on Biological Diversity, 2011). The Nagoya protocol specifically mentions the fair and equitable sharing of both monetary and non-monetary benefits, but there is still a need to understand what these benefits are, who is benefiting and whom those benefits could be shared with (Carroll et al., 2021; Chinsembu & Chinsembu, 2020). In parallel, the CARE Principles for data have been proposed specifically for the case of Indigenous Peoples (Carroll et al., 2020). Similar to the Nagoya Protocol, these principles emphasise the concept of Collective Benefit, seeking to ensure equitability.

Given the global interest and policies surrounding access to and benefits from biodiversity—culminating in the inclusion of Target 13 in the Kunming-Montreal Global Biodiversity Framework, which aims to ‘Increase the Sharing of Benefits From Genetic Resources, Digital Sequence Information, and Traditional Knowledge’—it is surprising that so little quantitative research has been conducted on this topic. Here we use a data-driven approach that makes use of open data from Global Biodiversity Information Facility (GBIF) and the free, collaborative knowledge base Wikidata (Wikidata.org) for drawing conclusions about what has been collected, where the specimens are now, who were the people involved in collecting them, and what those collections were and are used for.

We focus on two UK Overseas Territories in the Caribbean, Montserrat and the Cayman Islands. While these islands are quite small and distinct, they both have experienced a rich collecting history and share a colonial history, similar to much of the Global South, particularly in their isolation from the metropole and other countries where information on their biodiversity is held.

In conducting this research, we aspire to illuminate and contribute to reducing the historical inequities stemming from a colonial past, seeking effective resolutions and data repatriation, facilitating an improvement in the relationships between collection-holding institutions and the places where those collections originated.

## Materials and Methods

### Primary use of specimens from two UKOTs

To investigate who collected specimens on the case study islands, when they were collected, and for what research purposes, we followed a three-step process: 1) disambiguating the collector names recorded on specimen labels and digitised as ‘recordedBy’ in DarwinCore on GBIF, 2) establishing unambiguous links between collectors, their specimens and their publications, and 3) analysing biographical data and research outputs from collectors on Wikidata using SPARQL queries, and calculating the duration of collectors’ stays on the islands based on specimen collection dates.

For stage one, data of biological specimens collected from Montserrat and the Cayman Islands were extracted from the GBIF in three separate downloads. Specimens were extracted with ‘basisOfRecord’: MaterialSample, PreservedSpecimen and FossilSpecimen; with ‘occurrenceStatus’: present; and with ‘Administrative area’ (gadm.org): MSR (Montserrat), CYM (Cayman Islands), or a rectangular polygon that included all terrestrial parts of the two UKOTs and a substantial amount of the coastal waters (GBIF.org, 2022a, 2022b, 2022c). Deduplication and further screening of the data resulted in 17,907 specimens (Fig. S1).

Eighty-two percent of the specimens from Cayman Islands and 97% from the Montserrat specimens had ‘recordedBy’, a Darwin Core term with free-text, completed. The individual collector names were compiled in a spreadsheet. Character strings in the ‘recordedBy’ field that detailed multiple collectors were separated into individual names. Entries not referring to people, such as expedition names, were excluded.

Each individual name string was disambiguated with the aim of linking it to a biographical entry in Wikidata (Wikidata.org). If a person could be identified and deemed sufficiently notable for inclusion in Wikidata but did not yet have an entry, a new Wikidata entry was created for them (e.g. <u>Q117485794</u>^1^). Where biographical data, such as dates of birth and death, were available, this information was added to the corresponding Wikidata entity. Disambiguation followed the principles outlined in (Groom et al., 2022).

For the next step, each collector that had been identified uniquely to a Wikidata entry was unambiguously linked to their publications using the Wikimedia Toolforge Author Disambiguator tool (https://author-disambiguator.toolforge.org/) and to their specimens using Bionomia (https://bionomia.net/). A Bionomia public claims file was downloaded from https://bionomia.net/downloads on 28 December 2022 (Bionomia, 2024) and filtered to include only occurrences that matched the gbifID values from the Montserrat and Cayman Islands datasets. The filtered Bionomia attribution data contained 4,135 records. The relevant attributions from the Bionomia dataset were integrated into a PostgreSQL database that was created to store and manage the specimen occurrence data. The Cayman Islands dataset and the Montserrat dataset were also imported into the database. A table was created to store the disambiguated gbifID, recordedBy values, Wikidata identifiers and collection dates (Groom & Meeus, 2024a).

Finally, bespoke SPARQL queries were written to extract the biographical and bibliographical data directly from Wikidata (Groom & Meeus, 2024a). To understand the length of the stay of collectors on the islands the specimen collection dates and the biographical data of each collector was reviewed. In simple cases, a person had a cluster of specimen collection dates and these were interpreted as a single trip with start and end dates based on the earliest and latest specimen date. Such trips could often be cross-referenced to publications of the collector related to their expedition. For some collectors there were multiple clusters of recording dates, with long periods of no specimens in between. These were interpreted as multiple trips and the date of the first and last specimen from each cluster being used to estimate the length of the individual trip.

### Post-digitisation research applications of specimens

Since 2016, GBIF has been tracking the citation of data mobilised through its platform. These citation records are accessible through the GBIF Literature API, which can be found at https://techdocs.gbif.org/en/openapi/v1/literature. This service allows users to track which specimens have contributed to specific research articles, providing key metadata such as the Open Access status, article topics, and the country affiliations of the authors.

On 3rd June 2024, we extracted citing literature from the following types: “JOURNAL,” “WORKING_PAPER,” “BOOK,” and “BOOK_SECTION.” The extraction focused on literature with a relevance of “GBIF_CITED” and “GBIF_USED,” and including only peer-reviewed publications. After extracting the relevant literature, we downloaded all the GBIF-cited downloads from the literature identified in the first step, excluding any downloads that exceed 10 million rows because the specimens from smaller areas, such as islands, can represent only a tiny fraction of these massive datasets.

For each downloaded file, key information, including the gbifID, year of collection, country code, GBIF download key, and the publication DOI, was saved to an output file. All of this was conducted within a Jupyter notebook (Kluyver et al., 2016) entitled “linkLitToGbifId” (Groom, 2024a). The rows of output related to Montserrat and the Cayman Islands (represented by the country codes MS and KY, respectively) were then fed into a second Jupyter notebook called “GbifLitAnalysis.” This notebook extracted additional details about the publications using the GBIF Literature API (Groom, 2024b). Finally, a map was created using QGIS (QGIS Development Team, 2023), with a Mollweide equal area projection, to visualise the affiliations of the authors of the publications.

In a second phase, thematic analysis was conducted on the GBIF-cited literature to explore the prevalence and interrelationship of various research topics. Using a list of unique DOIs, topics associated with each publication were extracted and analysed. A network graph was constructed to visualise the co-occurrence of topics within the literature to provide insights into the thematic structure of biodiversity research on the two UKOTs, highlighting key areas of focus and their interconnections.

## Results

### Primary use of specimens from two UKOTs

Historical specimens from Montserrat and the Cayman Islands are mainly held in the USA, Canada and the United Kingdom (Fig 1 A,B). In the USA there are large collections, such as the Museum of Comparative Zoology at Harvard University and the Smithsonian National Museum of Natural History in Washington, D.C.; in Canada the Royal Ontario Museum and in the United Kingdom, the Natural History Museum, London. Likewise, the people who collected in these islands were largely from the USA and the UK, based upon where they were born, worked or died (Fig. 1 C,D). For example, prolific collectors include Wilmot Wood Brown Jr. (<u>Q109754544</u>), William Randolph Taylor (<u>Q21389931</u>) and Chapman Grant (<u>Q1062746</u>) who collected in the Cayman Islands, and Hugh Howard Genoways (<u>Q21341302</u>), Julius Boos (<u>Q26712297</u>) and Alexander Emanuel Agassiz (<u>Q122968</u>) who collected in Montserrat.

**Figure 1.**
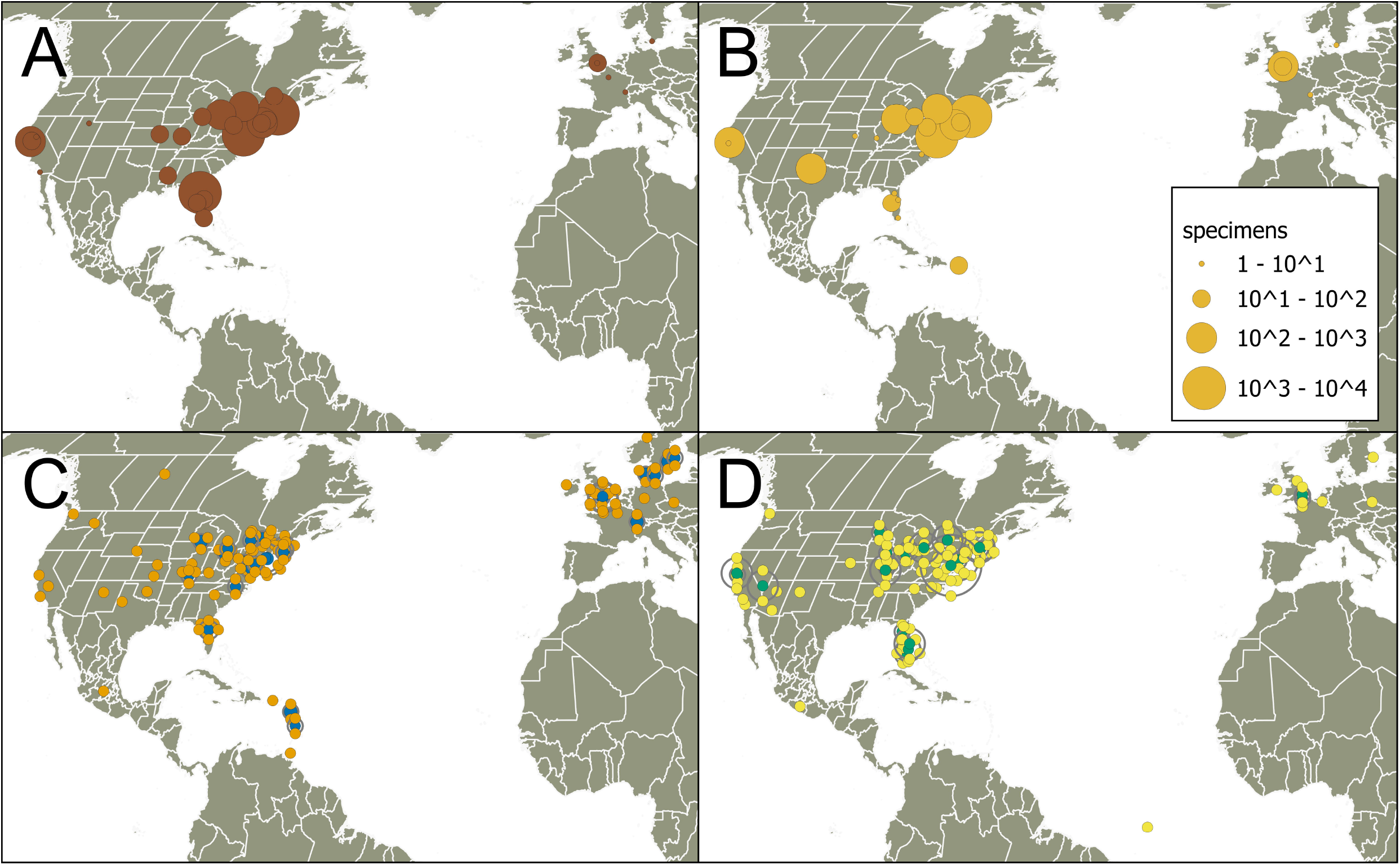
The institutions (A,B) and the places of birth, death, burial or employment (C,D) of collectors of specimens from Montserrat (A,C) and the Cayman Islands (B,D). Overlapping location data of the collectors has been displaced to ensure visibility. The data are taken from the biographies on Wikidata where those data exist.

We were able to disambiguate 113 of 198 (58%) names of collectors from Montserrat to entities in Wikidata, and 110 of 199 (55%) from the Cayman Islands. Twelve people collected on both islands. Four and eight people from the Cayman Islands and Montserrat respectively were identified as part of the local community. Their names are not in the public domain, except for on specimen labels, and therefore were not added to Wikidata. The disambiguated people contributed 10,622 specimens, 87% of attributed specimens from Montserrat and 63% from Cayman Islands. Those collectors were also authors of 1,586 and 2,465 scientific papers respectively.

The analysis of the biographies and publication records of the collectors shows that these people were mostly concerned with the documentation and description of the organisms of these islands. They can be described in the broad sense as taxonomists, though they vary in the taxonomic group they specialise in (Table S1 A,C,D). Montserrat perhaps attracts proportionally more botanists, but the Cayman Islands more ichthyologists (Table S1 C). The subjects of the papers they wrote show many on insects by Montserrat collectors and many on herpetology by collectors of the Cayman Islands (Table S1 A). They published in North American journals, and were often members of North American societies (Table S1 B,E). The data show no strong links with the United Kingdom, the metropole.

We also examined the duration of collecting trips based on the date of first and last specimen (Fig. S2). This task was made challenging, because there were many date errors in the digitised data. Such errors tended to mean collecting periods appeared longer than they were in reality. Nevertheless, 40% (MS, n=45) and 70% (KY, n=77) of collectors collected for two weeks or less on these islands.

While summary data give a panorama of the collector landscape, such a view misses the enormous heterogeneity of collectors. To give a clearer picture of the diversity of individual collectors we provide brief biographies of seven examples we selected for illustration (Groom & Meeus, 2024b). These profiles focus on the collectors’ visits to the islands, what they collected, the research they did, and where their collections are now.

### Post-digitisation research applications of specimens

At the time of the analysis (3rd June 2024), 6,273 papers that cite specimens in peer reviewed literature were downloaded from GBIF. This generated 312,820,974 specimens linked to 3,214 DOIs of downloads. A total of 190 publications were identified as citing or using Montserrat specimens, while 186 publications cited or used specimens from the Cayman Islands. Additionally, 126 publications referenced or used specimens from both islands. Due to the significant overlap in publications, the results were combined for analysis, resulting in a total of 250 publications. The papers date from 2016 to 2024, as it has only recently become possible to cite GBIF downloads using a DOI. Of these papers, 52% were Open Access. Although some of the cited specimens date back to the 1600s, the majority—85%—are from the 20th and 21st centuries.

None of the papers that used or cited specimens from the islands were researched specifically on the islands. In fact, the specimens from the islands only constitute a small proportion of specimens in all studies, even the publication with the highest use of species used less than 3% of specimens from the islands and averaged less than 0.1% of specimens. None of the authors of these papers were from the islands, though authors are widespread across the world, and although the USA is the most common origin of authors (13.5%), Brazil (8.6%), United Kingdom (7.2%), Germany (6.2) and China (6.0%) are also well represented (Fig. 2).

**Figure 2.**
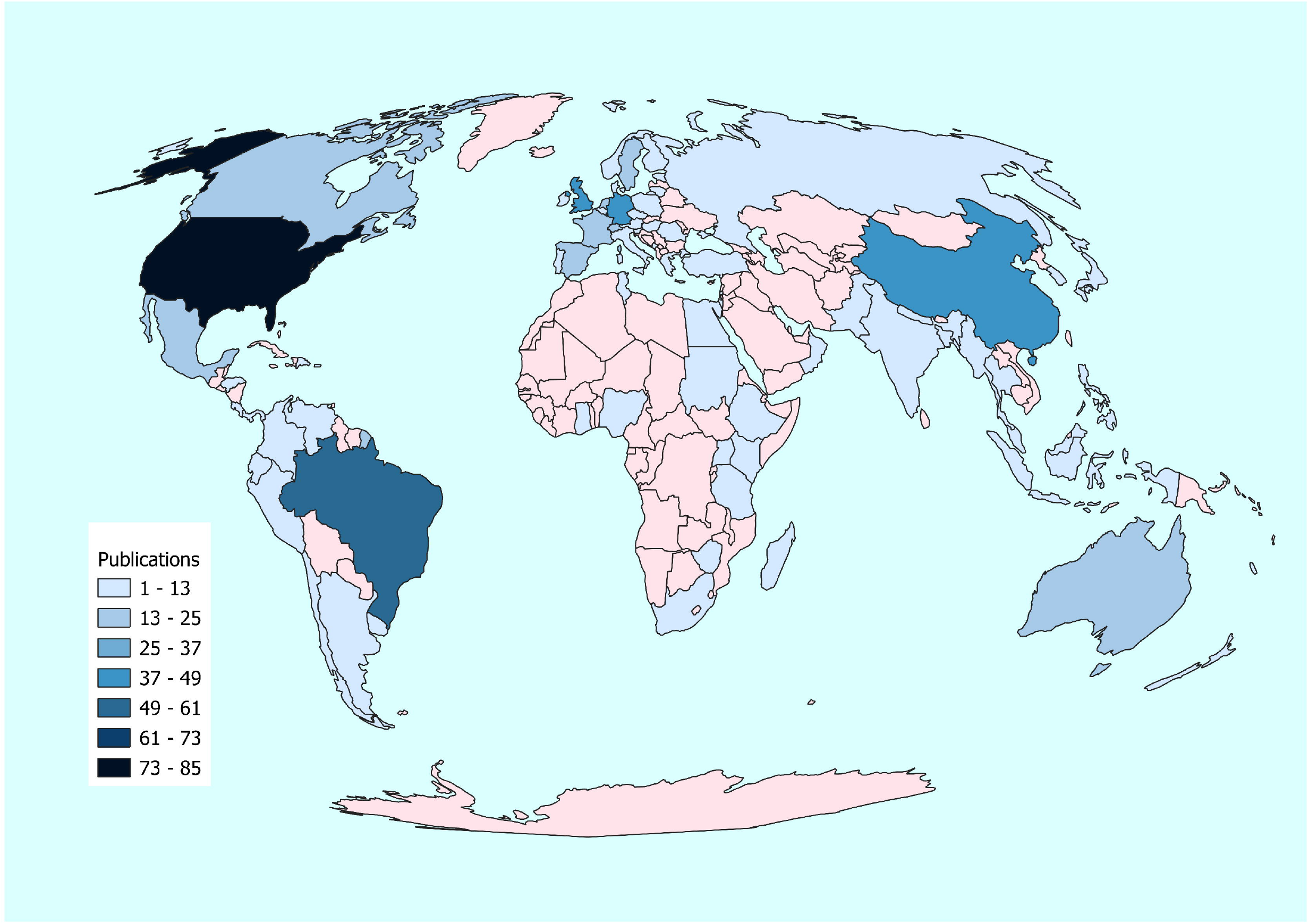
The origin of authors on publications using specimens from Montserrat and the Cayman Islands. The map uses an equal area Mollweide projection.

The network analysis of topics from publications citing specimens from Montserrat or the Cayman Islands reveals that “Ecology” is the most prominent theme, frequently co-occurring with topics like “Climate Change,” “Invasives,” and “Biogeography” (Fig. 3). Other significant topics include “Evolution” and “Species Distributions.” The network structure highlights strong connections between ecological and environmental themes, emphasising a particular focus on “Marine” and “Invasives”, with less emphasis on areas like “Taxonomy” and “Conservation”. This indicates that research involving specimens from these islands predominantly centres on evolutionary and biogeographical studies.

**Figure 3.**
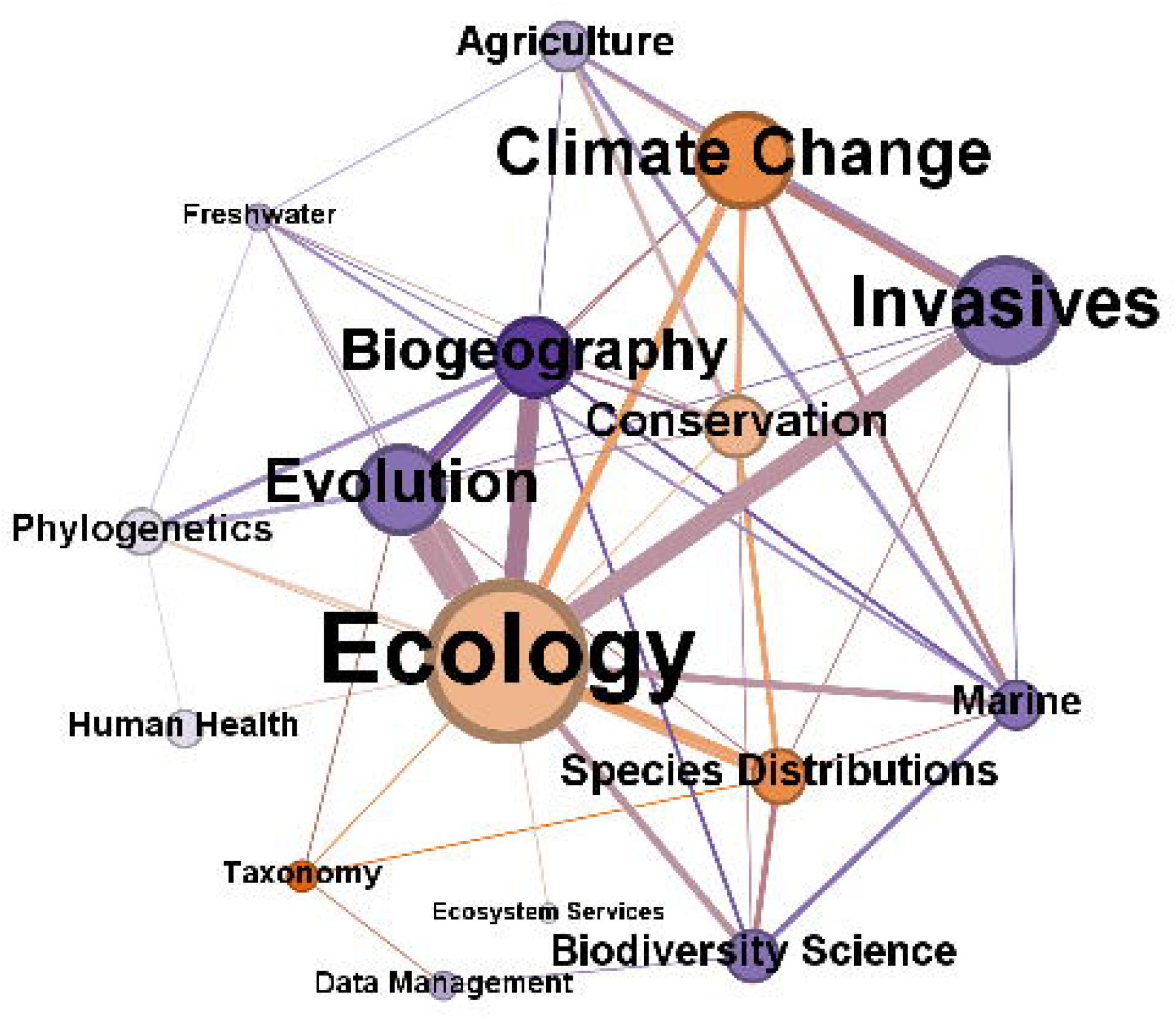
Topics associated with publications using digitised specimens from Montserrat or the Cayman Islands. The nodes represent the topics associated with each publication that cites or uses specimens from Montserrat or the Cayman Islands. Node size indicates the frequency of topic usage. Edges represent the co-occurrence frequency of topics within the same publication. Node colour reflects a comparison of the relative frequency of topic usage in the entire corpus of publications versus those specifically citing specimens from Montserrat or the Cayman Islands. Purple nodes indicate topics that are overused in comparison to publications from all countries, and orange nodes, underused.

## Discussion

This study provides insights into the history of biological collecting on Montserrat and the Cayman Islands in the Caribbean by examining the specimens, the collectors who gathered them, and the publications linked to these collections. Our analysis is based entirely on digitally available data, which introduces some biases. For example, we initially expected to see a greater presence of collectors from the UK, given its colonial history and extensive global collections. However, our results showed a higher representation of American collectors. This discrepancy may be due to slower progress in digitising and sharing UK collections compared to North American institutions, which benefit from initiatives like iDigBio (Nelson & Paul, 2019). Additionally, the geographic proximity of the USA likely contributes to a stronger presence of North American collectors.

As of October 2024, GBIF hosts over 251 million preserved specimen records. With global estimates of specimens in collections ranging from 1.2 to 2.1 billion (Ariño, 2010), this represents about 12% to 21% coverage. Since our initial GBIF data download approximately 1.5 years ago, the number of available specimens from Montserrat and the Cayman Islands has increased by roughly 1,000, reflecting ongoing digitization efforts. Major contributions have come from institutions like the Museum of Comparative Zoology at Harvard University and the Academy of Natural Sciences of Philadelphia. The continued digitisation of the Kew Herbarium will soon expand access to botanical specimens from the UK Overseas Territories, making even more records available on GBIF.

Despite our efforts to identify specimen collectors, many remain largely anonymous, even though their contributions have advanced scientific knowledge. This is often because they lack a digital presence, and we recognize that some may have preferred to remain unacknowledged or had no incentive to publicly disclose their work. Although these biases exist in the digital data, we believe there is sufficient information to draw meaningful conclusions, especially for collections in North America and Europe, where digitisation is more advanced.

With more than half of the collectors spending two weeks or less on the islands one could assume rampant ‘parachute science’, yet this would be a mischaracterization of some of it. Our results show that specimens from islands like Montserrat and Cayman Islands are relatively frequently used in studies of biogeography and evolution. The geological history of the Caribbean makes it a natural laboratory for evolution and this has obviously attracted such research (Mohammed et al., 2022 and references therein). Nonetheless, scientists can still share non-monetary benefits from their work, even during brief visits (Eichhorn et al., 2020). For example, researchers can present their findings on-site, exchange knowledge about species surveying techniques and species identification with local partners, ensure Open Access publication of their papers, openly publish underlying data, curate specimens for easy future use by local stakeholders, and public engagement (Edwards, 2004; Wilson et al., 2016; Park et al., 2023). When local collections are present, as they are on Montserrat and the Cayman Islands, visiting scientists can also contribute duplicate specimens to those collections, but this can only happen if such infrastructure is available. Short-term collecting does not have to be exploitative; avoiding the ‘parachute science’ label requires humility, respect for local biodiversity and knowledge, and inclusive practices.

While colonial patterns persist in some museums (Cisneros et al., 2022), others collection-holding institutes are working to reassess their roles in a postcolonial society (Antonelli, 2020; Das & Lowe, 2018; Gelsthorpe, 2021). Most research conducted by museums focuses on biodiversity documentation, conservation, biogeography, and phylogenetics—fields that fall under non-commercial research and do not generate revenue from intellectual property rights. Despite the benefits of such research, the sharing of these benefits has often been overlooked, especially in some national implementations of the Nagoya Protocol (Chinsembu & Chinsembu, 2020; Colella et al., 2023). Focusing solely on monetary benefits risks neglecting broader, non-monetary contributions to local communities. Examining specimens and collectors’ biographies reveals the substantial contributions of external research to the islands, including comprehensive biodiversity documentation, conservation studies, Floras and Faunas, and digital data infrastructure development, achievements unlikely to have been accomplished by the islands alone. Local administrations acknowledge this by facilitating research permits and welcoming visiting scientists.

However, there was little evidence during our studies, of collaboration of collectors with people on the islands. Islanders are sometimes mentioned as helpers (e.g. Corry et al., 2010 p.13), but rarely as co-authors (e.g. Ogrodowczyk et al., 2006; Dalsgaard et al., 2007) or co-collectors (GBIF.org, 2024). While the islands are too small to support major research institutions, local communities possess valuable knowledge and insights on species behaviour, population dynamics, genetics, and other studies involving local variability. For researchers, the pressure to publish for funding and career advancement is a significant driver, whereas non-scientists may not see authoring a paper as a worthwhile reward compared to everyday responsibilities. There is an opportunity for funders, educators, museums, and researchers to explore creative ways to foster collaboration and disseminate research outcomes more widely.

For example, in the Hidden Histories program funded by the UK Arts and Humanities Research Council (AHRC) and the Natural Environment Research Council (NERC), that funded this study, the funders allowed inclusion of partners from the Caribbean and those partners led part of the work and decided what the focus of the project would be. Not only did this provide an opportunity to integrate traditional knowledge into the research but it made it more equal and equitable. However, there was a cap on the maximum funding that non-academic partners could receive, and the principal investigator had to be based in a UK institution, eligible for UK Research and Innovation funding.

As part of this project, we developed a framework for best practices in environmental and other research with a focus on the UK Overseas Territories (Pienkowski & Wensink, 2022), complementing existing guides for research in the Global South (Haelewaters et al., 2021), and specific fields like field ecology (Baker et al., 2019) and biogeography (Eichhorn et al., 2020). This framework addresses different research stakeholders, such as funders, decision-makers, and researchers. Research funders vary significantly in how they structure funding schemes, including whether local partners can be included in grants and whether they have control over research priorities. Our best practices framework addresses these priorities for funding and advocates a more integrated approach to funding that better reflects the complexities and potential inequities in apportioning costs.

Dislocated specimens and inaccessible data can hinder local conservation efforts (Asase et al., 2022; Nakamura et al., 2024). While digitising natural history collections is a key step toward equitable access, it does not automatically ensure equitable use. We found that digitisation of collections largely contributes to research in the Global North, though it is encouraging that upper-middle-income countries, such as Brazil and China, have also been able to leverage this additional access. By tracking the use of digitised specimens, as we did here with the GBIF literature API, collection holders can assess whether the benefits from specimens are being equitably shared. To maximise the collective benefits of specimens, global institutions and local partners need to foster international collaborations and ensure mobilised data from natural history collections is being used and effectively applied to local, evidence-based conservation. This is essential for driving meaningful progress and promoting equitable knowledge sharing.

## Conclusions

The Caribbean is an exciting place for research, with its remarkable biological and geological diversity. Encouraging and supporting scientific studies in the region is crucial for advancing fundamental knowledge in biology and for addressing urgent conservation needs. The Caribbean’s biodiversity faces numerous threats, including climate change, sea-level rise, land-use changes, invasive non-native species, tourism, resource exploitation, and eutrophication. Museums and academic institutions can significantly contribute to these research efforts, but doing so effectively requires meaningful partnerships with the people from these islands. Collaborating with local partners not only respects their role as custodians of their biodiversity and traditional knowledge but also maximises the benefits for science, local communities, and conservation efforts.

Initially, we anticipated a stronger presence of UK collectors in our results, given the region’s colonial history and the vast amount of data from around the world housed in UK institutions. However, our findings showed otherwise, likely due to slower progress in digitising and sharing data from key institutions like the Royal Botanic Gardens, Kew; the Natural History Museum, London; and the herbaria at Cambridge, Oxford, and Manchester. While there are ongoing digitisation efforts, there is still significant room for improvement (Smith et al., 2022). Institutions and their funders should prioritise accelerating this process and actively consult with stakeholders in the territories to determine which collections should be prioritised for data sharing.

Research conducted on the islands has often been somewhat sporadic, with little overall coordination. This lack of systematic planning may have limited opportunities for local people to engage with, contribute to, or shape the research being conducted. While the creative freedom of scientists is essential for scientific progress, it is important to recognize that access to the natural environments where this research takes place is not an unlimited or freely available resource. Local communities possess extensive knowledge and offer a long-term perspective that is easily overlooked during brief research visits. To ensure that scientific work is truly inclusive and impactful, it must actively integrate local insights and priorities.

## Acknowledgements

The authors gratefully acknowledge the support of the Arts & Humanities Research Council and the Natural Environment Research Council under grant number AH/W008998/1. We also extend our thanks to our local partners for their assistance in clarifying information about collectors, as well as to the residents of Montserrat who participated in a workshop on data gathering, access, storage, and sharing to uncover their capacity needs.

We are grateful to Wikimedians Andra Waagmeester for assistance with SPARQL code to query Wikidata, and Sabine Von Mering for her guidance in setting up a Wikiproject. Additionally, we thank the GBIF technical staff for providing a comprehensive list of download keys along with their corresponding row counts.

Finally, we acknowledge the financial support provided by the Leopold III Fund and the Research Foundation – Flanders (FWO) for the field trip to QG and SM.

## Author Contribution

QG: Conceptualization, Data curation, Formal analysis, Investigation, Methodology, Project administration, Software, Validation,Visualization, Writing – original draft; SM: Data curation, Validation, Writing – review & editing; SB: Data curation; EC: Validation; SF: Funding acquisition, Validation; AJ: Funding acquisition; RM: Funding acquisition; MP: Funding acquisition; DP: Funding acquisition, Validation; CW: Funding acquisition; JP: Conceptualization, Funding acquisition, Project administration. All authors contributed to interpretations and revisions of the manuscripts.

## Data Availability Statement

The data that support the findings of this study are openly available in Zenodo at http://doi.org/10.5281/zenodo.13902532, http://doi.org/10.5281/zenodo.14056058, https://doi.org/10.5281/zenodo.13825234, https://doi.org/10.5281/zenodo.13823446.

The data on collectors are available in Wikidata at Wikidata.org, grouped under the WikiProject ‘Collectors of specimens from Montserrat and Cayman Islands’ (Q130465632). A comprehensive (dynamic) list with links to individual Wikidata records can be found here: https://www.wikidata.org/wiki/Special:WhatLinksHere/Q130465632. This data was retrieved from publicly available resources, backed by a reliable source and made available through Wikidata i.e. the public domain.

## Footnotes

1 John Kingsley Howes (1922-2013) was born in Montserrat and is referred to as Underwood’s collector in Corry et al. (2010). He collected the holotype of the Montserrat Galliwasp (*Diploglossus montserrati)*, a critically endangered endemic species of lizard described by Garth Underwood, a British herpetologist (Underwood, 1964; Stewart & Underwood, 2003) which is currently hosted in the Museum of Comparative Zoology at Harvard University. J. Kingsley Howes was part of the Formation Committee of the Montserrat National Trust (Montserrat Legislative Council, 1969) and was the manager of the Trants Estate Montserrat in 1986 (Pulsipher & Goodwin, 2001).

