## Supporting Information for "Capacity building needed to reap the benefits of access to biodiversity collections"

### *Plants, People, Planet* Supporting Information

The following Supporting Information is available for this article:

**Fig. S1** Data filtering process for specimen records from Montserrat and the Cayman Islands, inspired by the PRISMA method. The process included the identification of initial datasets from GBIF.org, removal of duplicates, and screening for specimens with accurate country codes. The final dataset consisted of 3,820 records from Montserrat and 14,087 records from the Cayman Islands.

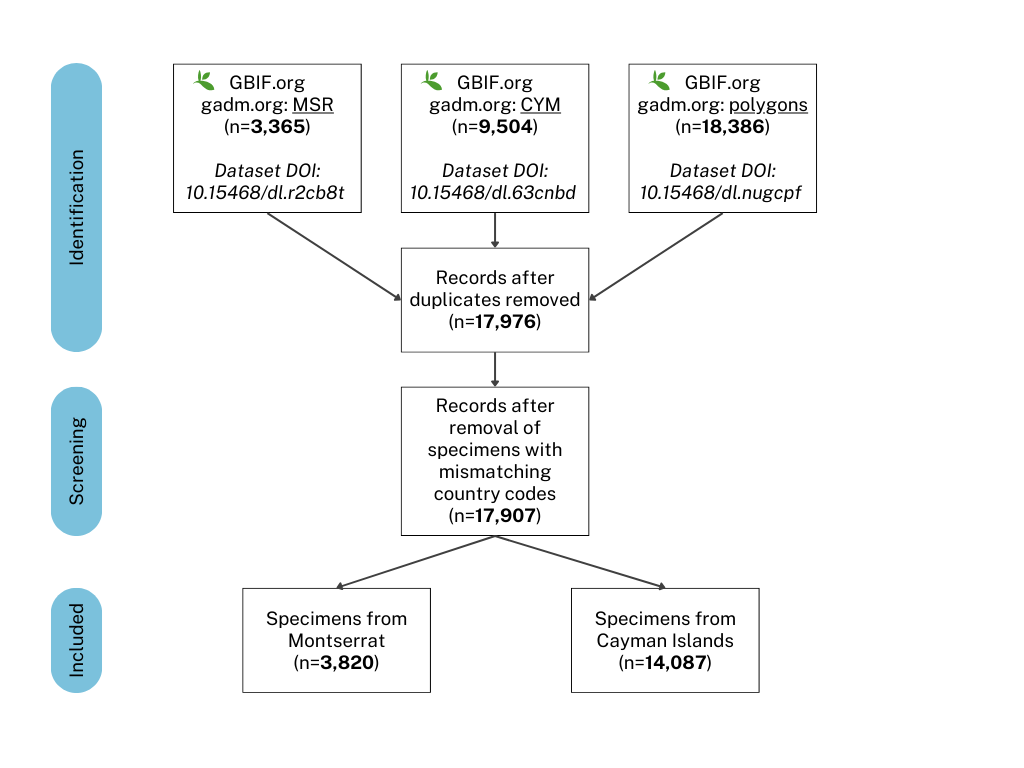

**Fig. S2** A time series of specimens collected in Cayman Islands and Montserrat

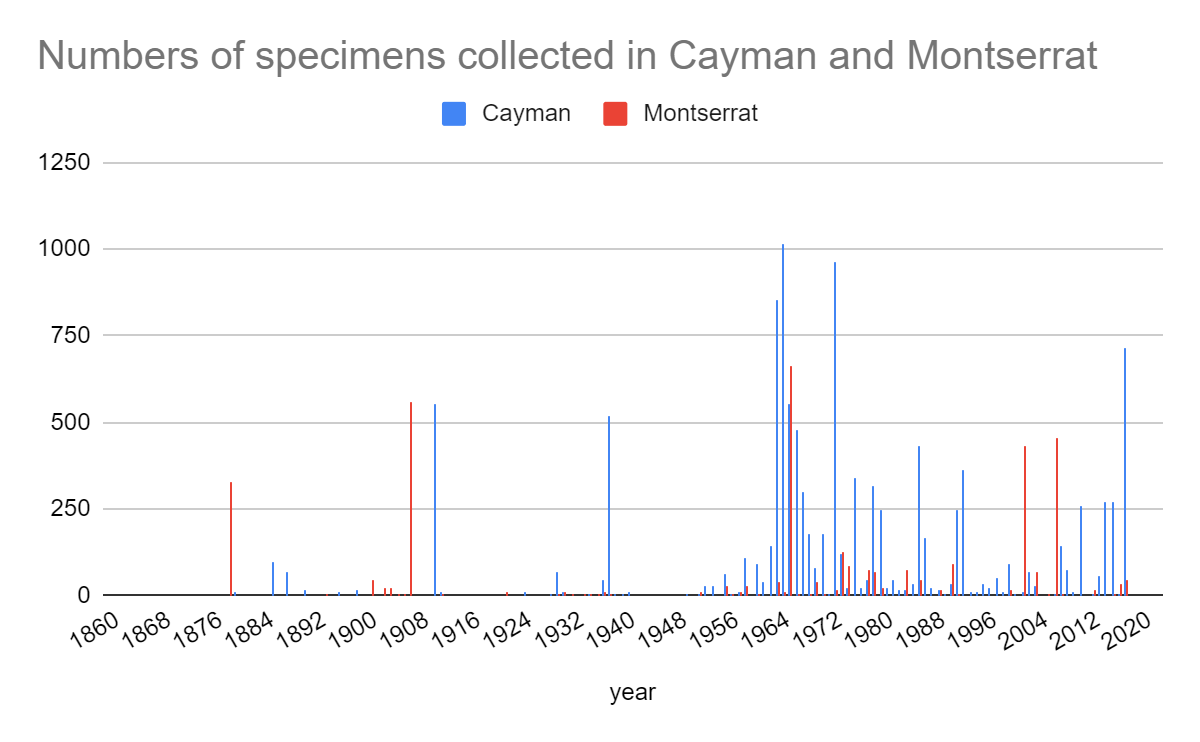

**Table S1** Characteristics of the collectors of Montserrat and the Cayman Islands. Results based on the collectors’ publication record.

|  | Montserrat | Cayman Islands |
| --- | --- | --- |
| A. What subjects do collectors write papers on? | beetles 72  species nova 68  Hymenoptera 63  Formicidae 46  Schistosoma mansoni 34  Orthoptera 28  Diptera 24  Biomphalaria 23  Crustacea 21  Lepidoptera 18 | species nova 231  Squamata 76  Euphorbiaceae 71  phylogenetics 65  Papua New Guinea 64  Sauria 54  Pharyngodonidae 45  Hymenoptera 44  Teleostei 41  Malpighiaceae 41 |
| B. Where were the articles published? | Proceedings of the United States National Museum 80 American Journal of Science 68  Mammalian Species 58  Journal of the Arnold Arboretum 56  The Florida Entomologist 52  The Coleopterists Bulletin 52  Occasional Papers, Museum of Texas Tech University 48  Journal of the Washington Academy of Sciences 40  Bulletin of the Torrey Botanical Club 35  Annals and Magazine of Natural History 32 | Copeia 241  Proceedings of the Academy of Natural Sciences of Philadelphia 214  Journal of Parasitology 101  Herpetologica 79  Evolution 65  Proceedings of the United States National Museum 61  Acta Parasitologica 61  The Auk 54  Taxon 50  The Florida Entomologist 48 |
| C. What were the occupations of the collectors? | zoologist 25  botanist 22  entomologist 21  researcher 16  university teacher 10  biologist 9  ornithologist 7  herpetologist 6  curator 5  writer 5 | zoologist 22  botanist 18  researcher 15  biologist 11  herpetologist 11  entomologist 10  ichthyologist 10  ornithologist 8  naturalist 6  curator 6 |
| D. What field did the collectors work in? | botany 10  zoology 7  entomology 6  herpetology 3  biology 3  literature 2  geology 2  bryology 2 | botany 6  herpetology 5  malacology 3  ornithology 3  ichthyology 3  systematics 2  biodiversity 2  marine biology 2 |
| E. What organisations were the collectors members of? | American Academy of Arts and Sciences 5  National Academy of Sciences 4  American Association for the Advancement of Science 3  Royal Swedish Academy of Sciences 2  Royal Prussian Academy of Sciences 2  German Academy of Sciences Leopoldina 2  Bavarian Academy of Sciences and Humanities 2 | American Academy of Arts and Sciences 4  National Academy of Sciences 4  American Association for the Advancement of Science 2  Santa Barbara Society of Natural History 1  Entomological Society of Canada 1  Washington Academy of Sciences 1  American Genetic Association 1  American Philatelic Society 1 |
